# Diversity chromosome evolution of Ty1-copia Retrotransposons in *Pennisetum purpureum* Revealed by FISH

**DOI:** 10.1101/2022.04.17.488575

**Authors:** Zehuai Yu, Yongji Huang, Xueting Li, Jiayun Wu, Muqing Zhang, Zuhu Deng

## Abstract

*Pennisetum purpureum* is a potential species for biofuel production. Characterization and chromosomal distribution of retrotransposons could enhance the comprehension of the role and dynamics of the repetitive elements in plants. In this study, a phylogenetic tree was constructed according to the conserved reverse transcriptase sequences and revealed that these Ty1-*copia* retrotransposons had typical structure. Analysis showed that total Ty1-*copia* retrotransposons had a significant component, as high as 5.12 × 10^3^ copy numbers in *P. purpureum*. Then, the chromosomal pattern of four known lineages were also analyzed with *Pennisetum glaucum* genome, which suggested that the Sire/Maximus lineage had the highest copy number and followed by Tork/Angela, Tork/TAR, Retrofit/Ale. Additionally, the chromosomal distribution of total Ty1-*copia* retrotransposons was detected by fluorescence *in situ* hybridization (FISH) to be a dispersed pattern with weak clustering, mostly near the centromeric regions of *P. purpureum* chromosomes; interestingly, there were four obvious signals in the subterminal chromosomes. These results suggested that there occurred differential dynamic evolution directions of Ty1-copia retrotransposons within *P. purpureum*. Furthermore, co-localization of Ty1-*copia*, 5S rDNA, and 35S rDNA indicated that two chromosome 2 and four chromosome 4 were identified. Concurrently, subterminal signals of Ty1-*copia*-type retrotransposons were located on four other homologous chromosomes. Altogether, these results shed light on the diversification of Ty1-*copia* retrotransposons and have the significance for generation of valid chromosomal markers in retrotransposon families.

## Introduction

Developing a new conversion technology has recently attracted increased attention for its potential value for producing bioethanol and methane (Howard et al. 2003, Menegol et al. 2014). The tropical species *P. purpureum* represents a new alternative energy crop that may provide abundant and sustainable lignocellulosic biomass which can be used for biofuel production (Xie et al. 2011, Li et al. 2013). As a C4 plant, elephant grass can suppress photorespiration and has great potential to efficiently convert solar energy to biomass (Zhu et al. 2008). This grass also has high productivity compared to other species, producing a large productions per year (Woodard and Prine 1993, Somerville et al. 2010). In addition, this biomass can be grown on different soil types, requiring little additional nutrients for growth. Due to these characteristics, elephant grass is a favorable option as a second-generation ethanol production.

*P. purpureum* is a tetraploid species (2n = 4*x* = 28) that belongs to an economically important tropical forage plant in the genus *Pennisetum*. The DNA content per haploid nucleus for *P. purpureum* was estimated by flow cytometry to be approximately 1.15 pg (1.12×10 ^3^ Mb) (Martel et al. 1997). Repetitive sequences account for a considerable part of the plant genome, which constitute up to 80% in some plants (Vicient et al. 2001). These motifs can be divided into two broad groups as follows: tandemly repeated and dispersed sequences that include transposable elements (Heslop-Harrison 2000). Transposable, or mobile, elements are divided into two main classes, DNA transposons and retrotransposons (Wicker et al. 2007). In general, retrotransposons include long terminal repeats (LTRs) or non-LTR retrotransposons. The content of retrotransposons leads to large disparity genome in different plants. Retrotransposons ranged from 10% ∼ 70% in rice (Chen et al. 2013, Kawahara et al. 2013), maize (Schnable et al. 2009) and sorghum (Paterson et al. 2009), resulting in different genomes. Moreover, different LTR retrotransposons in plants result in various chromosomal patterns. LTR retrotransposons are mostly concentrated in highly heterochromatic regions (centromeres, pericentromeres, telomeres) (Cheng et al. 2002, Devos et al. 2002, Jiang et al. 2002, Ammiraju et al. 2007, Gao et al. 2009) and have a tendency to accumulate in the pericentromeric regions of the genome. While, Xu and Du found that LTR retrotransposons are distributed in euchromatic regions in *Solanum lycopersicum* (Xu and Du 2014).

LTR retrotransposons include Ty1-*copia*, Ty3-gypsy, Bel-Pao, retroviruses, and endogenous retroviruses (Kumar and Bennetzen 1999, Kumar and Bennetzen 2000)(Kumar and Bennetzen 1999, Kumar and Bennetzen 2000), and of these, Ty1-*copia* and Ty3-gypsy retrotransposons are abundant in plant genomes (Wicker et al. 2007, de la Chaux and Wagner 2011). In Ty1-*copia* elements, there have four subunits including protease, integrase, reverse transcriptase (RT), and ribonuclease H. To date, although genome sequencing can greatly facilitate the analysis of retrotransposons, the cost and assembly of repetitive sequences make it difficult in non-model plant or genomic complexity species. Hence, amplifying Ty1-*copia* RT domains is more feasible and has led to a better understanding of sequence evolution and phylogenetic relationship in numerous plants (Flavell 1992, Pearce et al. 1996, Goodwin and Poulter 2002, Santini et al. 2002, Ma et al. 2008, Atefeh et al. 2013, Kolano et al. 2013, Li et al. 2013). Additionally, the content of Ty1-*copia* elements is indispensable to study the dynamics of genome size in plant. The relative content of Ty1-*copia* RT sequences in plant genomes has been widely assessed by dot-blot hybridization (Ma et al. 2008, Kolano et al. 2013). The chromosomal pattern of Ty1-*copia* RT sequence is also vital for studying the retrotransposition dynamics of their retrotransposons. FISH is an efficient molecular technique for karyotype analysis, chromosome recombination, and chromosomal transmission in plant genomes (Abd Eltwab 2003, Wu et al. 2014, Han et al. 2015). 35S rDNA and 5S rDNA sequences are “hot” markers widely used to analyze chromosome evolution and karyotype for FISH (D’Hont et al. 1998, Zhang et al. 2016). Several studies have shown that Ty1-*copia* sequences are ubiquitous in the plant genome, just like 35S rDNA and 5S rDNA markers. However, there are no studies on the relationship between localization of Ty1-*copia* RT, 35S rDNA, and 5S rDNA in *P. purpureum*.

In this study, we isolated the conserved Ty1-*copia* RTs from the *P. purpureum* genome to identify retrotransposon sequences and evaluated their heterogeneity, relationship, abundance, and chromosomal patterns. Furthermore, we also analyzed the genomic distribution of Ty1-*copia* RT, 35S rDNA, and 5S rDNA in *P. purpureum*.

## Materials and methods

### Plant materials and DNA isolation

Fujian elephant grass (*P. purpureum*, 2*n* = 4*x* = 28) were provided by the Sugarcane Research Institute of Yunnan Agriculture Science Academy. Young leaves were collected from the greenhouse at Fujian Agriculture and Forestry University in Fuzhou, China. A standard cetyltrimethyl ammonium bromide (CTAB) protocol was used for extracting total genomic DNA (gDNA) (Stewart and Via 1993).

### PCR amplification and cloning of PCR products

The RT region of the Ty1-*copia* retrotransposons was amplified by PCR using gDNA of *P. purpureum* as templates and a pair of degenerate primers (Ty1-F: 5’-ACNGCNTT(C/T)(C/T)TNCA(C/T)GG-3’; Ty1-R: 5’-A(A/G)CAT(A/G)TC(A/G)TCNAC(A/G)TA-3’) (Flavell 1992). PCR amplification was performed on a Veriti^@^96-well Thermal Cycler (Applied Biosystems, United States). Reaction mixtures were 50 μL, containing 50 ng of gDNA, 20 pmol of each primer. Reactions were denatured at 94°C for 4 min, followed by 35 cycles of 94°C denaturation for 50 s, annealing at 45°C for 45 s, extension at 72°C for 30 s, and final extension at 72°C for 5 min. Then, to avoid PCR preference, we selected three times PCR production mixtures for further cloning and sequencing. In total, 88 Ty1-*copia* RT sequences were named as PpTy1-*copia*-1 to PpTy1-*copia*-88 and available in the GenBank database MH674202-MH674289.

### Sequence analysis, comparisons, and phylogenetic trees

Identified Ty1-*copia* RT sequences from *P. purpureum* were compared with other Ty1-*copia* RT that come from graminaceous species (*Saccharum, Sorghum, Erianthus, Triticum, Zea, Oryza*, and *Hordeum*). MEGA 7 software was used to align these nucleotide sequences (Kumar et al. 2016). These amino acid sequences were obtained from Ty1-*copia* RT sequences by the EMBOSS package2 (https://www.ebi.ac.uk/Tools/st/emboss_transeq/). MACSE was used to align Ty1-*copia* RT amino acid sequences that used in this study (Ranwez et al. 2011). Multiple sequence alignment of the Ty1-*copia* RT amino acid sequences was undertaken by using MUSCLE (Santos et al. 2008). MEGA 7 was used to constructed phylogenetic tree using neighbor-joining method (Kumar et al. 2016). TBtools software was used for sequences local blast with *P. glaucum* genome (Chen et al. 2018), and these positions were visualized with Integrative Genomics Viewer software (Thorvaldsdóttir et al. 2012).

### Dot-blot hybridization

Reverse dot-blot hybridization (RDB) was performed to detect the content of Ty1-*copia* RT sequences in *P. purpureum* genome. All purified plasmids were quantified in NanoVue PlusTM (GE Healthcare, Princeton, NJ, United States) and a final concentration of 50 ng/μL plasmids was used for hybridization. The *P. purpureum* gDNA probe was labeled with digoxigenin-11-dUTP (DIG) using a DIG Nick Translation Kit (Roche Diagnostics). Hybridization was performed as described in the Instruction Manual of the DIG High Prime DNA Labeling and Detection Starter Kit I (Roche Diagnostics). Washing conditions was performed according to Huang et al (Huang et al. 2017).

The copy number of total Ty1-*copia* RT sequences in *P. purpureum* was estimated by dot-blotting. Serial dilutions of *P. purpureum* gDNA (500, 400, 250, 200, 100, and 50 ng) and total Ty1-*copia* RT plasmids (2, 1.5, 1, 0.75, 0.5, 0.375, and 0.25 ng) were used further hybridization. Total Ty1-*copia* sequences labeled with DIG were prepared as probes using the PCR-DIG Probe Synthesis Kit (Roche). ImageJ and Calculator software (http://cels.uri.edu/gsc/cndna.html) were conducted to estimate the copy number per genome of total Ty1-*copia* RT (Schneider et al. 2012).

### FISH

Meristem of the *P. purpureum* root-tips were prepared according to Huang et al (Huang et al. 2017). Total Ty1-*copia* RT sequences of *P. purpureum* was labeled with DIG using PCR as described in the PCR-DIG Probe Synthesis Kit (Roche Diagnostics). 35S rDNA was labeled with DIG or biotin-16-dUTP (Roche) using a Nick Translation Kit (Roche Diagnostics). 5S rDNA labeled with biotin-16-dUTP (Roche) using the PCR-DIG Probe Synthesis Kit (Roche Diagnostics). FISH was performed as described by Huang et al (Huang et al. 2017). Chromosomes were counterstained with DAPI (4′, 6-diamidino-2-phenylindole, 6 μg/mL). An AxioScope A1 Imager fluorescent microscope (Carl Zeiss, Gottingen, Germany) was used for capturing image.

## Results

### PCR products and characterization of Ty1-*copia* RT sequences

The PCR amplification yielded a product of the expected size (approximately 260 bp) in *P. purpureum*. 88 independent clones were randomly selected clones from three times PCR production mixture to avoid PCR biased. 84 sequences (95.5 %) ranged from 253 to 266 in length and four sequences (4.5 %) ranged from 315 to 316 bp. All translated nucleotide sequences had TAFLHG, S/ALYGLKQ, and YVDDM conserved domains. There were four longer Ty1-*copia* RT sequences that varied from 315 to 316 bp (PpTy1-*copia* 39, 74, 78, and 79) and differed from the other plants but shared the upstream TAFLHG, the central conserved domain S/ALYGLKQ, and downstream YVDDM motifs. However, all longer Ty1-*copia* RT sequences were defective (Table S1). In the other 84 Ty1-*copia* RT sequences, 20 sequences were defective (Table S1). In total, 27.27 % of Ty1-*copia* RT sequences may be non-functional in *P. purpureum*.

### Classification and phylogenetic analysis of identified Ty1-*copia* RT sequences from *P. purpureum*

All identified 88 Ty1-*copia* RT nucleotide sequences were translated to 74-95 amino acids in length. Very high sequence heterogeneity was existed among these amino acid sequences, and they could be classified into 28 distinct groups (Fig 1). These groups were then nominated as Pc (*P. purpureum copia*-type retrotransposon) and named from Pc1 to Pc28. These groups had different amino acid similarity, ranging from 61.8% to 100% (Table S1).

**Fig 1.**
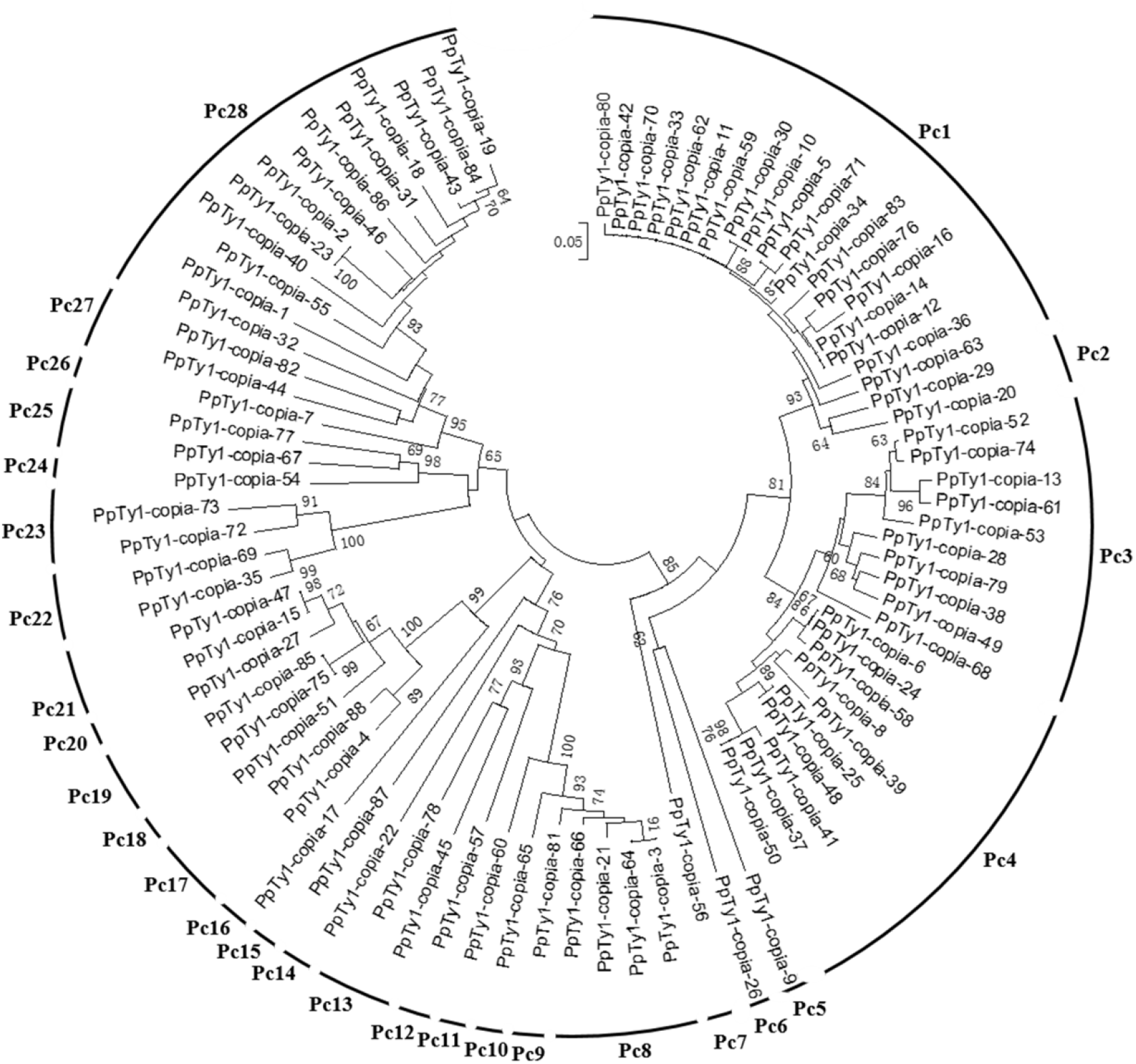
Phylogenetic analysis of Ty1-*copia* RT isolated from *P. purpureum*.

A neighbor-joining tree was constructed by aligning Ty1-*copia* RT amino acid sequences among the named Pc1-28 Ty1-*copia* RT sequences with related Ty1-*copia* retrotransposons from other graminaceous species (Fig 2). These were separated into six distinct evolutionary lineages (I–VI) of four known lineages (Tork/TAR, Tork/Angela, Sire/Maximus, and Retrofit/Ale) and two unclassified families. Lineage I contained Pc8-Pc13, which was clustered with a Tork/TAR lineage of *Triticum, Saccharum, Oryza*, and *Zea*, indicating that the Tork/TAR lineage have the largest known groups of Ty1-*copia* in *P. purpureum* (Fig 2). Among the Ty1-copia RT sequences, 4 (40 %) occurred frameshifts or stop codons; whereas the remaining 6 (60 %) contained potentially functional Ty1-copia RT domains. Lineage II contained Pc14-15, which was closed to a Tork/Angela lineage of *Oryza* and *Hordeum*, respectively. While, all of these putative amino acids were functional in *P. purpureum*. Lineage III contained Pc22-Pc23, which was clustered with a Sire/Maximus lineage of *Saccharum*. In this lineage, all of the Ty1-copia RT sequences included stop coding or frameshift mutations. Lineage IV contained Pc24-Pc28, which was clustered with a Retrofit/Ale lineage of *Saccharum, Triticum, Oryza, Sorghum*, or *Zea*. This lineage included the most numerous clones (19 elements) in known lineages, and of these, 14 (73.7%) putative amino acids contained potentially functional Ty1-copia RT domains. These results indicated that RT sequences might share the common origin and that horizontal transmission of retrotransposons has occurred among the gramineous plants. In especial, all these groups were closed to *Saccharum*, suggesting that *P. purpureum* have potential to be use for sugarcane breeding to broaden its genetic base.

**Fig 2.**
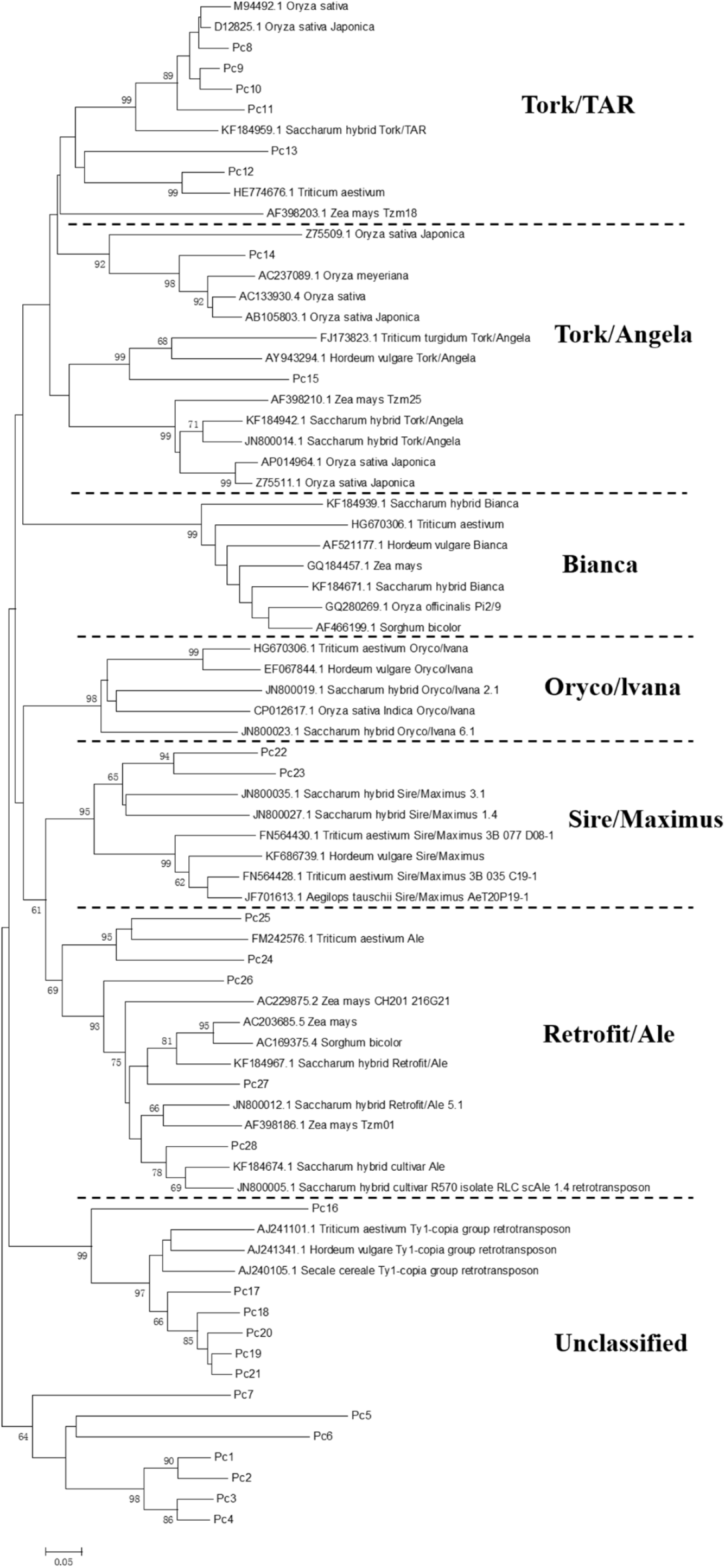
Phylogenetic analysis of amino acid sequences based on Ty1-*copia* RT sequences from *P. purpureum* and other graminaceous species (*Saccharum, Oryza, Hordeum, Sorghum, Triticum*, and *Zea*). Bootstrap values over 60 are indicated at the nodes.

However, the other groups clustered to unclassified lineage. Lineage V contained Pc16-Pc21, which was clustered with an unclassified lineage but had high homology to Ty1-*copia*-like elements belonging to other graminaceous species, including *Triticum, Hordeum*, and *Secale*. The last lineage contained Pc1-Pc7, which was clustered with an independent branch in the evolutionary tree. Altogether, these results demonstrated homology between *P. purpureum* and other gramineous plants, but also indicated certain heterogeneity (Fig 2).

### Relative abundance in *P. purpureum* genome and chromosomal patterns of RTs

RDB was performed to estimate the relative content of Ty1-*copia* RT from the *P. purpureum* genome. All purified plasmids of Ty1-*copia* RT sequences were used to hybridize to the genomic DNA of *P. purpureum*. While, there were no obvious hybridization signals, suggesting that single Ty1-*copia* RT sequence had a low content in *P. purpureum* genome. Then, Total Ty1-*copia* sequences were used to hybridize to the genomic DNA of *P. purpureum*. Results showed that they produced obvious hybridization signals, confirming that total Ty1-*copia* retrotransposons had a large content in the *P. purpureum* genome (Fig 3). A quantitative dot-blot assay was performed to detect the copy number of the total Ty1-*copia* RT in the *P. purpureum* genome, using serial dilutions of total Ty1-*copia* RT as probes and genomic DNA from *P. purpureum*. To improve the accuracy of the detection, we set up three technical repeats. The hybridization intensity data indicated that the copy number of total Ty1-*copia* RT sequences was approximately 5.12 × 10^3^ in a haploid nucleus (Fig 3).

**Fig 3.**
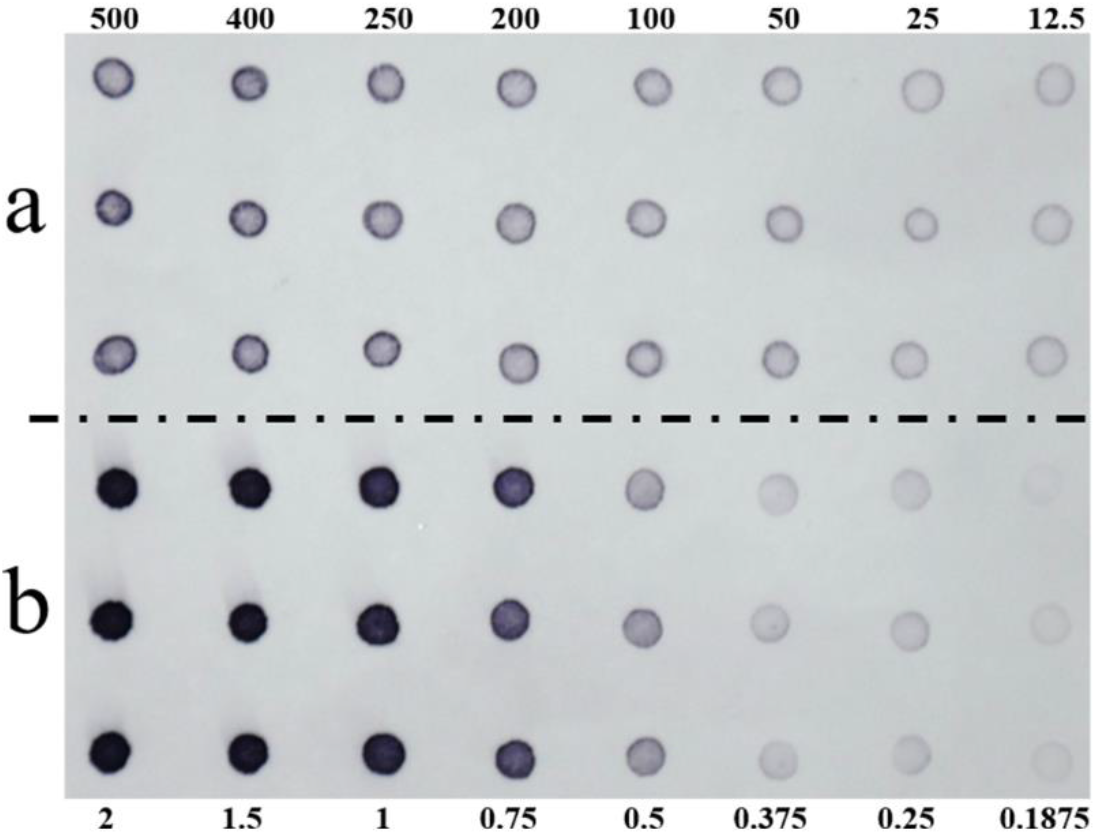
Estimation of total copy number of Ty1-*copia* RT sequences in *P. purpureum* genome. Serial dilutions (ng) of genomic DNA from *P. purpureum* (row a) and Ty1-*copia* RT sequences plasmids (row b).

FISH was carried out on interphase nuclei and somatic metaphase chromosomes for investigating the chromosomal pattern of Ty1-*copia* RT sequences in the *P. purpureum* genome. The DAPI-positive heterochromatic region was colored by Ty1-*copia* probes in interphase nuclei (Fig 4). However, feeble hybridization signals were observed on the euchromatic regions of interphase nuclei. Interestingly, the total Ty1-*copia* RT probes were differentially dispersed on somatic metaphase chromosomes. These signals showed a dispersed pattern with weak clustering, mostly distributed near the centromere region in somatic metaphase chromosomes (Fig 4); Furthermore, we also found that the total Ty1-*copia* retrotransposons had four obvious concentrated signals on interphase nuclei (Fig 4B). These signals were located exclusively in the distal regions of four chromosomes in metaphase chromosomes (Fig 4D). These results suggested that there were two different drove evolution of Ty1-*copia* retrotransposons in *P. purpureum*.

**Fig 4.**
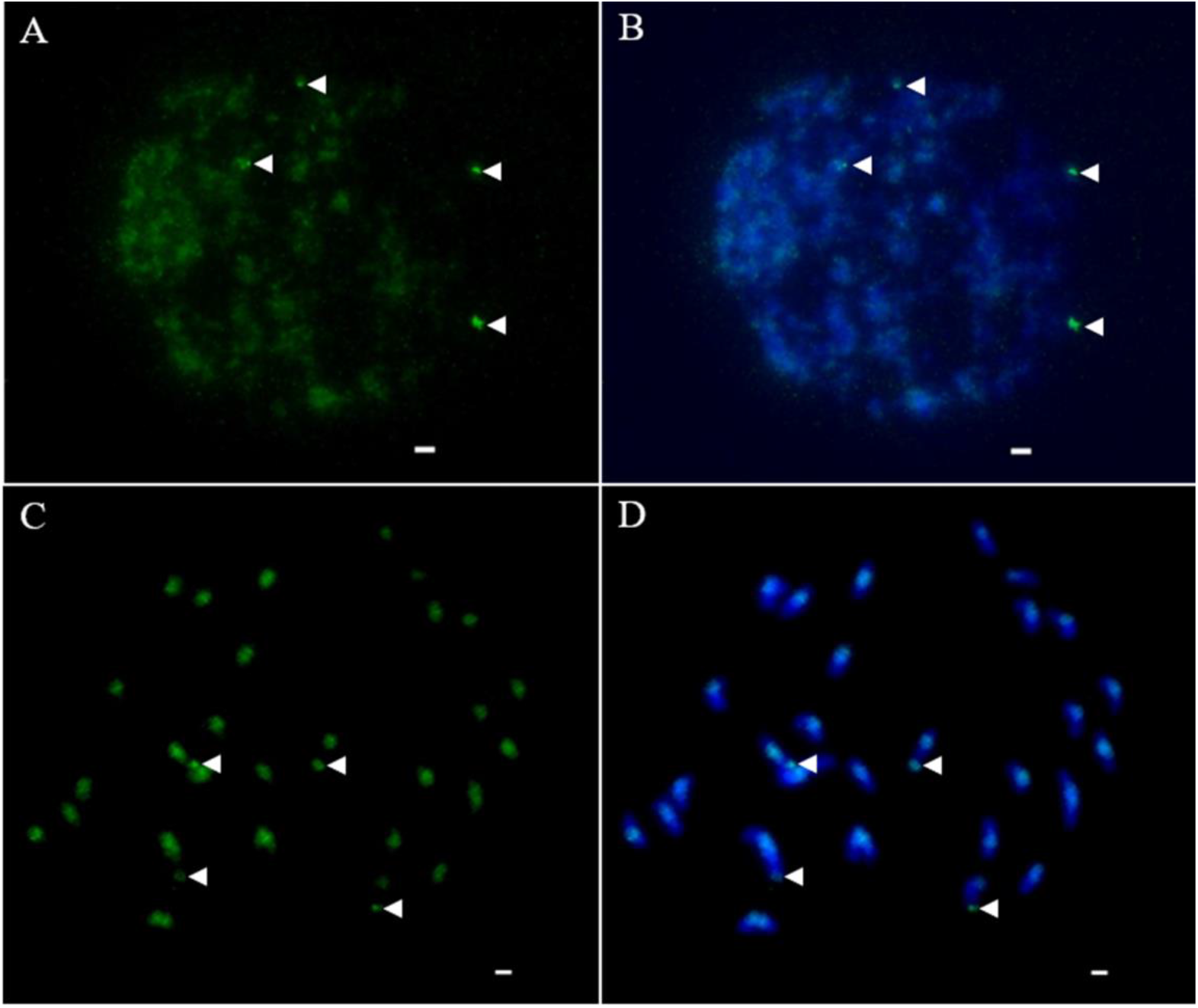
FISH results of Ty1-*copia* RT sequences on interphase nuclei and metaphase chromosomes of *P. purpureum* (2n = 4*x* = 28). Nucleus and metaphase chromosomes stained with DAPI (B and D). Total Ty1-*copia* RT probes on interphase nucleus (A) and metaphase chromosomes (C). White arrows indicated obvious concentrated signals in telomeres. Scale bars = 5 μm.

### Co-localization of Ty1-*copia*, 35S rDNA, and 5S rDNA in *P. purpureum*

Identifying the chromosomal locations of *copia*-type retrotransposons could facilitate the selection of families for informative chromosomal marker in karyotype. The previous paper reported that *P. purpureum* is an allotetraploid (A’A’BB) and confirmed that genome A’A’ has a high homology with *P. glaucum* (genome A, 2n = 2*x* = 14). Hence, the chromosome distribution of our identified four known Lineages, Tork/TAR, Tork/Angela, Sire/Maximus and Retrofit/Ale was evaluated using the published *P. glaucum* genome. Results suggested that the Sire/Maximus had 1052 copies in *P. glaucum*. The copy number of Tork/Angela, Tork/TAR and Retrofit/Ale were 551, 131 and 116 respectively (Fig 5). Notably, most Sire/Maximus sequences were closed to the centromeric regions in *P. glaucum* chromosomes (Fig 5), these results were partly consistent with the FISH location of RTs in *P. purpureum* (Fig 4). The conserved 5S and 35S rDNAs sequences were also detected in chromosome 2 and chromosome 4 (Fig 5).

**Fig 5.**
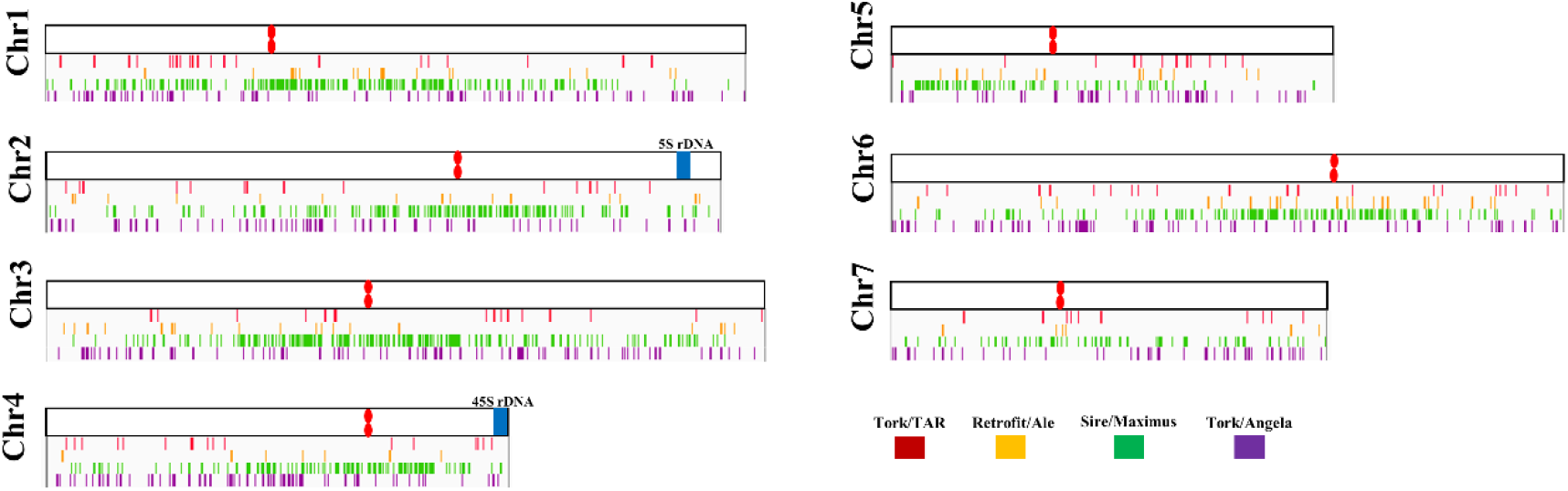
Chromosomal distributions of RTs, 35S rDNA and 5S rDNA in *P. glaucum* genome. PpTy-copia-21, PpTy-copia-46, PpTy-copia-69 and PpTy-copia-87 sequences represent the Tork/TAR, Tork/Angela, Sire/Maximus and Retrofit/Ale, and were used for locating in *P. glaucum* genome.

Further, the chromosomal location of 35S rDNA and 5S rDNA in the *P. purpureum* genome were also detected. FISH analysis was performed using digoxigenin (DIG)-labeled 35S rDNA and biotin-labeled 5S rDNA as probes. The 35S rDNA probe was located in the distal regions of four chromosome 4, region was distinctly decondensed compared to the rest of the chromosome (Fig 6). Two of the four homologous chromosomes lacked the 5S rDNA loci, and the 5S rDNA probe was only located in two chromosome 2 (Fig 6). In addition, a double-label FISH assay was performed using digoxigenin (DIG)-labeled Ty1-*copia* sequences, biotin-labeled 35S rDNA and 5S rDNA as probes to examine their co-localization in *P. purpureum*. As expected, *copia*-type retrotransposon was located in the distal regions of four chromosomes homologous chromosomes (Fig 7). Moreover, these probes were found to be non-overlapping, indicating that the *copia*-type retrotransposons may be a reliable marker for chromosome identification and karyotyping studies in *P. purpureum* (Fig 7D).

**Fig 6.**
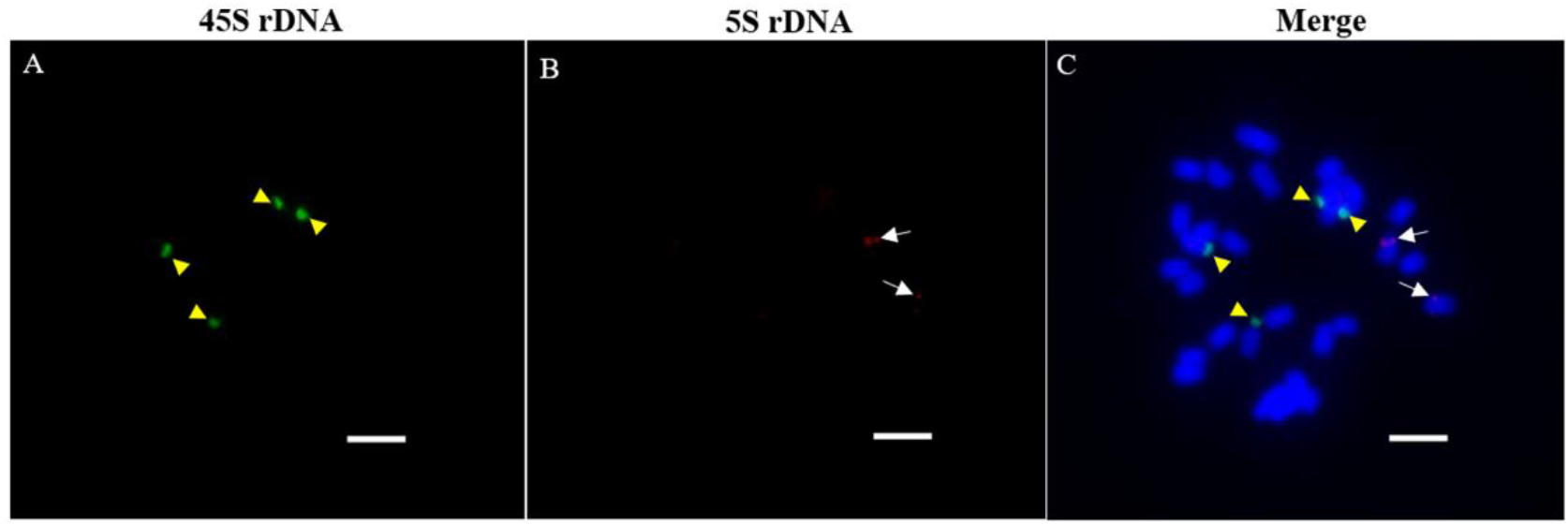
Localization of 35S rDNA and 5S rDNA probes on root-tip metaphase chromosomes of *P. purpureum* (2n = 4*x* = 28) by FISH. Yellow arrows indicated 35S rDNA signals (green). Long white arrows indicated 5S rDNA signals (red). Scale bars = 5 μm.

**Fig 7.**
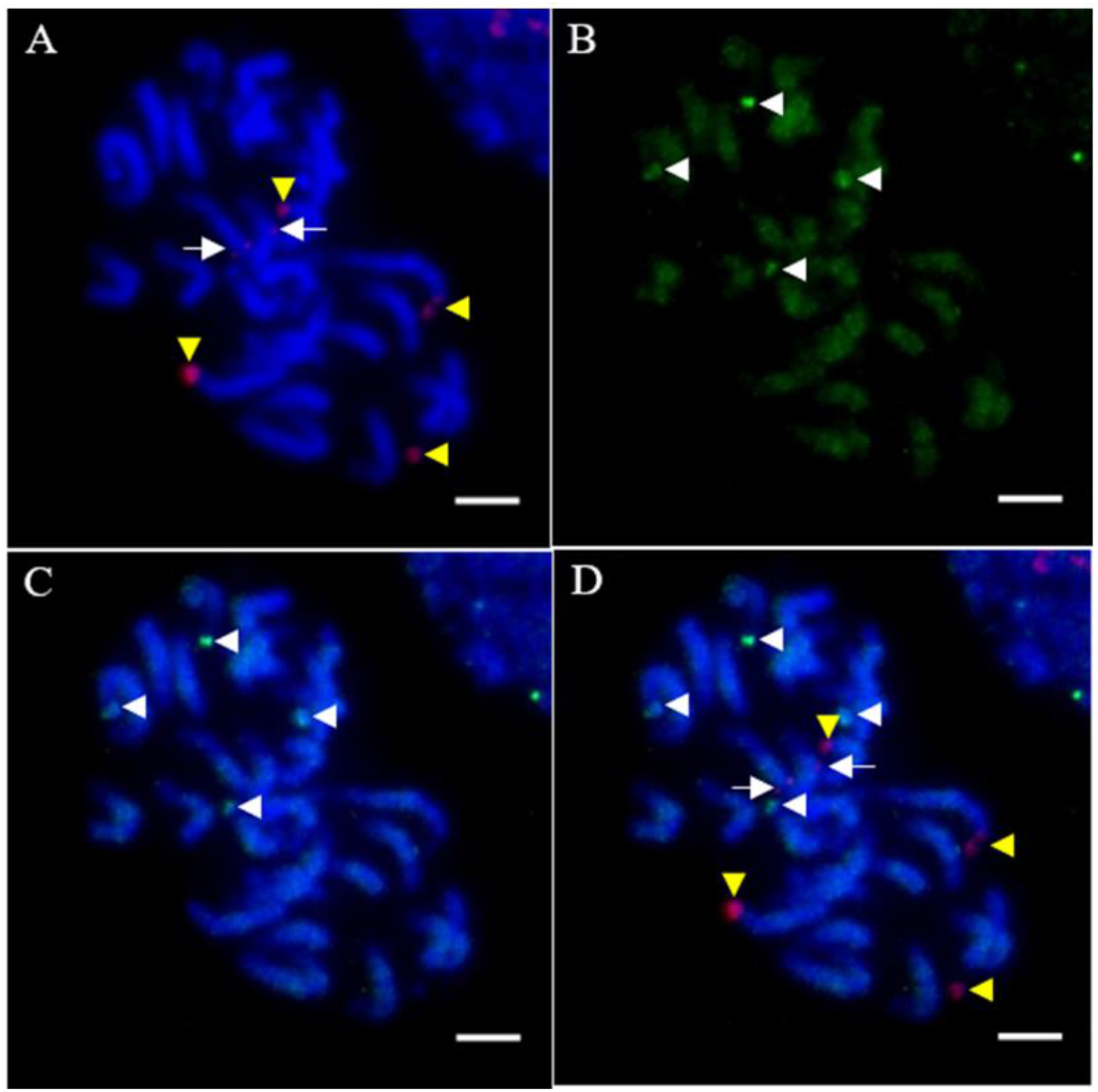
Localization of Ty1-*copia* 35S rDNA and 5S rDNA probes on root-tip metaphase chromosomes of *P. purpureum* (2n = 4*x* = 28) by FISH. White arrows indicated Ty1-*copia* signals (green). Yellow arrows indicated 35S rDNA signals (red). Long white arrows indicated 5S rDNA signals (red). Scale bars = 5 μm.

## Discussion

*P. purpureum* belongs to an economically important tropical forage plant in the genus *Pennisetum*. Although it is becoming an increasingly popular crop that can be used to produce clean energy ethanol, its allotetraploid genome origin, structure, and evolution have not yet been thoroughly studied. Retrotransposons will drive genetic diversity that can potentially cause alters in genome structure and gene expression, and these elements are thought to be crucial for genome plasticity and evolution (Kidwell and Lisch 1997, J et al. 2005, Hawkins et al. 2006, Zedek et al. 2010, D 2013). Owing to genome sequencing is limited in non-model plant, using PCR to amplify conserved regions of Ty1-*copia* retrotransposons has resulted in explosive number of molecular level researches of these elements on evolution and phylogenetic relationships in different plants (Flavell 1992, Pearce et al. 1996, Goodwin and Poulter 2002, Santini et al. 2002, Ma et al. 2008, Atefeh et al. 2013, Kolano et al. 2013, Lee et al. 2013, Huang et al. 2017).

In this study, a high degree of heterogeneity was be found in Ty1-*copia* retrotransposon sequences from the *P. purpureum*, consistent with other previous reported in various plants (Goodwin and Poulter 2002, Santini et al. 2002, Ma et al. 2008, Atefeh et al. 2013, Kolano et al. 2013, Lee et al. 2013, Huang et al. 2017). Results indicated that these Ty1-copia RT sequences clustered into six main subgroups in a highly ramified tree, indicating the large genetic variability within this species. Reis et al. showed that *P. purpureum* is an allotetraploid (A’A’BB) and confirmed that genome A’A’ has a high homology with *P. glaucum* (genome A, 2n = 2*x* = 14), while distinguished from the 14 chromosomes from genome BB (Reis et al. 2014). Therefore, such large genetic variability of Ty1-*copia* retrotransposon may result from different genomes in *P. purpureum*, as some belong to the A’ genome while others belong to the B genome. Out of 88 clones, 27.27 % of Ty1-*copia* RT sequences may be non-functional in *P. purpureum*, owing to appear stop codons or frameshifts; of these, all longer Ty1-*copia* RT sequences included stop codons. These results suggested that the heterogeneous elements may derived from the accumulation of mutations of defective elements in the host genome. This phenomenon is also ubiquitous in other plants, with a defective rate as high as >80 % (Huang et al. 2017). Additionally, Pc1-Pc7 lineages were an independent clade in the evolutionary tree in graminaceous species, indicating that these sequences have potential to be a PCR molecular marker to assist the plant breeding in *P. purpureum* in the future.

The relative content of the Ty1-copia retrotransposons were nearly 5.12 × 10^3^ per genome in *P. purpureum*, indicating that Ty1-*copia* retrotransposons play a vital role in the genome evolution of *P. purpureum*. Retrotransposons make a great contribution to the evolution of plant genome size, as retrotransposons comprise less than ∼5%-17% in a small genome, while as high as ∼70%-75% in a large genome (Vicient et al. 1999, McCarthy et al. 2002, Pereira 2004, Schnable et al. 2009). In this study, total Ty1-*copia* sequences showed dispersed organization across all chromosomes. Similarly, intrachromosomal localization was revealed in many taxonomic groups plant species (Pearce et al. 1996, Atefeh et al. 2013). Although total Ty1-*copia* sequences showed dispersed organization, stronger signals emerged near the centromere region (Fig 3). Usually, pericentromeric regions contain fewer genes and may show suppressed genetic recombination. Hence, it will result in the enrichment of Ty1-*copia* retrotransposons in the centromere region. Interestingly, there were other four homologous chromosomes have an obvious clustering in telomeres, indicating the presence of Ty1-*copia* sequences in chromosomal distal regions with high copies (Fig 3). There were two followed possible that Ty1-*copia* RT inserted near telomeres: either the element prior inserts into subtelomeric regions or it inserts randomly throughout the genome; however, when it inserts into gene-rich regions, it would be eliminated by selection against deleterious effects of genes and chromosomal rearrangements. Altogether, differential evolution pattern of Ty1-*copia* RTs within this species suggested that *P. purpureum* will be an ideal plant for further speculate the molecular evolution determinants of Ty1-*copia* sequences, pericentromeric or telomeric targeting.

Ty1-copia RT have a great potential to become a valid chromosomal marker as they were uniquely located in four chromosomes in *P. purpureum*. In this study, 35S rDNA and 5S rDNA probes were located in four chromosome 4 and two chromosome 2, respectively. This distribution pattern of 35S rDNA and 5S rDNA loci is consistent with previous studies, indicating that two 5S rDNA loci were lost or too weak to detect (Zhao 2010). Co-localization of Ty1-*copia*, 35S rDNA, and 5S rDNA showed that these markers were exclusively located in different non-homologous chromosomes, making Ty1-*copia* sequences suitable to identify chromosome in *P. purpureum*. The physical pattern of retrotransposons will specifically broaden the way to label narrow chromosomal regions or particular sub-genome, as demonstrated in the previous reports (Wang et al. 1999, Hanson et al. 2000). The location of Ty1-*copia* RT will be helpful for further studying the potential interspecific gene flow between a crop and its related species, and obtaining valid information about recombination and introgression.

## Conclusion

This is the first molecular cytological evidence of phylogenetic diversity, genomic abundance, and chromosomal pattern of Ty1-*copia* retrotransposons in *P. purpureum*. We isolated and characterized Ty1-*copia* retrotransposons, which were then clustered four known lineages (Tork/TAR, Tork/Angela, Sire/Maximus, and Retrofit/Ale) and two unclassified lineages. These retrotransposons are approximately 5.12 × 10^3^ per haploid genome in the *P. purpureum*. FISH results showed that total Ty1-copia sequences had differential distributions, pericentromere or subtelomere region. Furthermore, co-localization of Ty1-*copia*, 35S rDNA, and 5S rDNA confirmed that these sequences were located in different homologous chromosomes. Altogether, these results will be helpful for further understanding the dramatic evolution of Ty1-*copia* retrotransposons in *P. purpureum* and indicate an additional marker for identifying chromosome analysis in *P. purpureum*.

## Acknowledgements

We thank the Sugarcane Research Institute of Yunnan Agriculture Science Academy for providing the plant materials used in this study. We greatly appreciate Bioscience Editing Solutions for critically reading this paper and providing helpful suggestions.

## References

Abd Eltwab, M. H. (2003). “Physical mapping of 35S rDNA loci by fluorescent in situ hybridization and evolution among polyploid Dendranthema species.” Chromosome Sci 7(3): 71–76.

Ammiraju, J. S. S., A. Zuccolo, Y. Yu, X. Song, B. Piegu, F. Chevalier, J. G. Walling, J. Ma, J. Talag, D. S. Brar, P. J. SanMiguel, N. Jiang, S. A. Jackson, O. Panaud and R. A. Wing (2007). “Evolutionary dynamics of an ancient retrotransposon family provides insights into evolution of genome size in the genus Oryza.” Plant J 52(2): 342–351. doi: 10.1111/j.1365-313X.2007.03242.x.

Atefeh, A., T. Suguru, S. Hiroe, O. Nobuko and F. Kiichi (2013). “Structural characterization ofcopia-type retrotransposons leads to insights into the marker development in a biofuel crop, Jatropha curcas L.” Biotechnol Biofuels 6(1): 1–13.

Chen, C., R. Xia, H. Chen and Y. He (2018). “TBtools, a Toolkit for Biologists integrating various HTS-data handling tools with a user-friendly interface.” bioRxiv: 289660.doi: 10.1101/289660.

Chen, J., Q. Huang, D. Gao, J. Wang, Y. Lang, T. Liu, B. Li, Z. Bai, J. Luis Goicoechea, C. Liang, C. Chen, W. Zhang, S. Sun, Y. Liao, X. Zhang, L. Yang, C. Song, M. Wang, J. Shi, G. Liu, J. Liu, H. Zhou, W. Zhou, Q. Yu, N. An, Y. Chen, Q. Cai, B. Wang, B. Liu, J. Min, Y. Huang, H. Wu, Z. Li, Y. Zhang, Y. Yin, W. Song, J. Jiang, S. A. Jackson, R. A. Wing, J. Wang and M. Chen (2013). “Whole-genome sequencing of Oryza brachyantha reveals mechanisms underlying Oryza genome evolution.” Nat Commun 4(4): 1595.doi: 10.1038/ncomms2596.

Cheng, Z., F. Dong, T. Langdon, S. Ouyang, C. R. Buell, M. Gu, F. R. Blattner and J. Jiang (2002). “Functional rice centromeres are marked by a satellite repeat and a centromere-specific retrotransposon.” Plant Cell 14(8): 1691–1704.

D’Hont, A., D. Ison, K. Alix, C. Roux and J. C. Glaszmann (1998). “Determination of basic chromosome numbers in the genus Saccharum by physical mapping of ribosomal RNA genes.” Genome 41(2): 221–225.

D, L. (2013). “How important are transposons for plant evolution?” Nature Reviews. Genetics 14(1): 49–61.

de la Chaux, N. and A. Wagner (2011). “BEL/Pao retrotransposons in metazoan genomes.” BMC Evol Biol 11(1): 1–16.doi: Artn 15410.1186/1471-2148-11-154.

Devos, K. M., J. K. M. Brown and J. L. Bennetzen (2002). “Genome size reduction through illegitimate recombination counteracts genome expansion in Arabidopsis.” Genome Res 12(7): 1075–1079.doi: 10.1101/gr.132102.

Flavell, A. J. (1992). “Ty1-copia group retrotransposons and the evolution of retroelements in the eukaryotes.” Genetica 86(1-3): 203–214.

Gao, D., N. Gill, H. R. Kim, J. G. Walling, W. Zhang, C. Fan, Y. Yu, J. Ma, P. Sanmiguel and N. Jiang (2009). A lineage-specific centromere retrotransposon in Oryza brachyantha.

Goodwin, T. J. and R. T. Poulter (2002). “A group of deuterostome Ty3/ gypsy-like retrotransposons with Ty1/ copia-like pol-domain orders.” Mol Genet Genomics 267(4): 481–491.doi: 10.1007/s00438-002-0679-0.

Han, Y. H., T. Zhang, P. Thammapichai, Y. Q. Weng and J. M. Jiang (2015). “Chromosome-Specific Painting in Cucumis Species Using Bulked Oligonucleotides.” Genetics 200(3): 771–779.doi: 10.1534/genetics.115.177642.

Hanson, R. E., M. N. Islam-Faridi, C. F. Crane, M. S. Zwick, D. G. Czeschin, J. F. Wendel, T. D. McKnight, H. J. Price and D. M. Stelly (2000). “Ty1-copia-retrotransposon behavior in a polyploid cotton.” Chromosome Res 8(1): 73–76.

Hawkins, J. S., H. Kim, J. D. Nason, R. A. Wing and J. F. Wendel (2006). “Differential lineage-specific amplification of transposable elements is responsible for genome size variation in Gossypium.” Genome Res 16(10): 1252–1261.doi: 10.1101/gr.5282906.

Heslop-Harrison, J. S. (2000). “RNA, genes, genomes and chromosomes: repetitive DNA sequences in plants.” Chromosomes 13: 45-+.

Howard, R. L., E. Abotsi, E. J. V. Rensburg and S. Howard (2003). “Lignocellulose biotechnology: issues of bioconversion and enzyme production.” Afr J Biotechnol 2: 602–619.

Huang, Y., L. Luo, X. Hu, F. Yu, Y. Yang, Z. Deng, J. Wu, R. Chen and M. Zhang (2017). “Characterization, Genomic Organization, Abundance, and Chromosomal Distribution of Ty1-copia Retrotransposons in Erianthus arundinaceus.” Front Plant Sci 8(924): 924.doi: 10.3389/fpls.2017.00924.

J, M. S. P, L. J M. J and B. Jl (2005). “DNA rearrangement in orthologous orp regions of the maize, rice and sorghum genomes.” Genetics 170(3): 1209–1220.

Jiang, N., Z. Bao, S. Temnykh, Z. Cheng, J. Jiang, R. A. Wing, S. R. McCouch and S. R. Wessler (2002). “Dasheng: A recently amplified nonautonomous long terminal repeat element that is a major component of pericentromeric regions in rice.” Genetics 161(3): 1293–1305.

Kawahara, Y., M. de la Bastide, J. P. Hamilton, H. Kanamori, W. R. McCombie, S. Ouyang, D. C. Schwartz, T. Tanaka, J. Wu, S. Zhou, K. L. Childs, R. M. Davidson, H. Lin, L. Quesada-Ocampo, B. Vaillancourt, H. Sakai, S. S. Lee, J. Kim, H. Numa, T. Itoh, C. R. Buell and T. Matsumoto (2013). “Improvement of the Oryza sativa Nipponbare reference genome using next generation sequence and optical map data.” Rice (N Y) 6(1): 4.doi: 10.1186/1939-8433-6-4.

Kidwell, M. G. and D. Lisch (1997). “Transposable elements as sources of variation in animals and plants.” P Natl Acad Sci USA 94(15): 7704–7711.doi: DOI 10.1073/pnas.94.15.7704.

Kolano, B., E. Bednara and H. Weiss-Schneeweiss (2013). “Isolation and characterization of reverse transcriptase fragments of LTR retrotransposons from the genome of Chenopodium quinoa (Amaranthaceae).” Plant Cell Rep 32(10): 1575–1588.doi: 10.1007/s00299-013-1468-4.

Kumar, A. and J. L. Bennetzen (1999). “Plant retrotransposons.” Annu Rev Genet 33(1): 479–532.doi: 10.1146/annurev.genet.33.1.479.

Kumar, A. and J. L. Bennetzen (2000). “Retrotransposons: central players in the structure, evolution and function of plant genomes.” Trends Plant Sci 5(12): 509–510.

Kumar, S., G. Stecher and K. Tamura (2016). “MEGA7: Molecular Evolutionary Genetics Analysis Version 7.0 for Bigger Datasets.” Mol Biol Evol 33(7): 1870–1874.doi: 10.1093/molbev/msw054.

Lee, S. I., K. C. Park, J. H. Son, Y. J. Hwang, K. B. Lim, Y. S. Song, J. H. Kim and N. S. Kim (2013). “Isolation and characterization of novel Ty1-copia-like retrotransposons from lily.” Genome 56(9): 495–503.doi: 10.1139/gen-2013-0088.

Li, H. Q., C. L. Li, T. Sang and J. Xu (2013). “Pretreatment on Miscanthus lutarioriparious by liquid hot water for efficient ethanol production.” Biotechnol Biofuels 6(1): 76.doi: 10.1186/1754-6834-6-76.

Ma, Y., H. Sun, G. Zhao, H. Dai, X. Gao, H. Li and Z. Zhang (2008). “Isolation and characterization of genomic retrotransposon sequences from octoploid strawberry (Fragaria x ananassa Duch.).” Plant Cell Rep 27(3): 499–507.doi: 10.1007/s00299-007-0476-7.

Martel, E., D. DeNay, S. SiljakYakovlev, S. Brown and A. Sarr (1997). “Genome size variation and basic chromosome number in pearl millet and fourteen related Pennisetum species.” J Hered 88(2): 139–143.doi: DOI 10.1093/oxfordjournals.jhered.a023072.

McCarthy, E. M., J. Liu, G. Lizhi and J. F. McDonald (2002). “Long terminal repeat retrotransposons of Oryza sativa.” Genome Biol 3(10): RESEARCH0053.

Menegol, D., A. Luisi Scholl, R. Claudete Fontana, A. J. Pinheiro Dillon and M. Camassola (2014). “Potential of a Penicillium echinulatum enzymatic complex produced in either submerged or solid-state cultures for enzymatic hydrolysis of elephant grass.” Fuel 133: 232–240.doi: https://doi.org/10.1016/j.fuel.2014.05.003.

Paterson, A. H., J. E. Bowers, R. Bruggmann, I. Dubchak, J. Grimwood, H. Gundlach, G. Haberer, U. Hellsten, T. Mitros, A. Poliakov, J. Schmutz, M. Spannagl, H. Tang, X. Wang, T. Wicker, A. K. Bharti, J. Chapman, F. A. Feltus, U. Gowik, I. V. Grigoriev, E. Lyons, C. A. Maher, M. Martis, A. Narechania, R. P. Otillar, B. W. Penning, A. A. Salamov, Y. Wang, L. Zhang, N. C. Carpita, M. Freeling, A. R. Gingle, C. T. Hash, B. Keller, P. Klein, S. Kresovich, M. C. McCann, R. Ming, D. G. Peterson, R. Mehboobur, D. Ware, P. Westhoff, K. F. Mayer, J. Messing and D. S. Rokhsar (2009). “The Sorghum bicolor genome and the diversification of grasses.” Nature 457(7229): 551–556.doi: 10.1038/nature07723.

Pearce, S. R., U. Pich, G. Harrison, A. J. Flavell, J. S. Heslop-Harrison, I. Schubert and A. Kumar (1996). “The Ty1-copia group retrotransposons of Allium cepa are distributed throughout the chromosomes but are enriched in the terminal heterochromatin.” Chromosome Res 4(5): 357–364.

Pearce, S. R., U. Pich, G. Harrison, A. J. Flavell, J. S. Heslopharrison, I. Schubert and A. Kumar (1996). “TheTy1-copia group retrotransposons ofAllium cepa are distributed throughout the chromosomes but are enriched in the terminal heterochromatin.” Chromosome Research An International Journal on the Molecular Supramolecular & Evolutionary Aspects of Chromosome Biology 4(5): 357–364.

Pereira, V. (2004). “Insertion bias and purifying selection of retrotransposons in the Arabidopsis thaliana genome.” Genome Biol 5(10): R79.doi: 10.1186/gb-2004-5-10-r79.

Ranwez, V., S. Harispe, F. Delsuc and E. J. Douzery (2011). “MACSE: Multiple Alignment of Coding SEquences accounting for frameshifts and stop codons.” PLoS One 6(9): e22594.doi: 10.1371/journal.pone.0022594.

Reis, G. B. D., A. T. Mesquita, G. A. Torres, L. F. Andrade-Vieira, A. V. Pereira and L. C. Davide (2014). “Genomic homeology between Pennisetum purpureum and Pennisetum glaucum (Poaceae).” Comp Cytogenet 8(3): 199–209.

Santini, S., A. Cavallini, L. Natali, S. Minelli, F. Maggini and P. G. Cionini (2002). “Ty1/copia-and Ty3/gypsy-like DNA sequences in Helianthus species.” Chromosoma 111(3): 192–200.doi: 10.1007/s00412-002-0196-2.

Santos, G., I. M. C. Maia, F. F. Araújo, A. Pacheco, S. Medeiros, C. R. F. Gadelha, S. C. Silva, R. Araujo-Filho and D. Oliveira (2008). HiperMUSCLE: An intuitive graphical user interface for the multiple sequence alignment program MUSCLE (Edgar, 2004).

Schnable, P. S., D. Ware, R. S. Fulton, J. C. Stein, F. Wei, S. Pasternak, C. Liang, J. Zhang, L. Fulton, T. A. Graves, P. Minx, A. D. Reily, L. Courtney, S. S. Kruchowski, C. Tomlinson, C. Strong, K. Delehaunty, C. Fronick, B. Courtney, S. M. Rock, E. Belter, F. Du, K. Kim, R. M. Abbott, M. Cotton, A. Levy, P. Marchetto, K. Ochoa, S. M. Jackson, B. Gillam, W. Chen, L. Yan, J. Higginbotham, M. Cardenas, J. Waligorski, E. Applebaum, L. Phelps, J. Falcone, K. Kanchi, T. Thane, A. Scimone, N. Thane, J. Henke, T. Wang, J. Ruppert, N. Shah, K. Rotter, J. Hodges, E. Ingenthron, M. Cordes, S. Kohlberg, J. Sgro, B. Delgado, K. Mead, A. Chinwalla, S. Leonard, K. Crouse, K. Collura, D. Kudrna, J. Currie, R. He, A. Angelova, S. Rajasekar, T. Mueller, R. Lomeli, G. Scara, A. Ko, K. Delaney, M. Wissotski, G. Lopez, D. Campos, M. Braidotti, E. Ashley, W. Golser, H. Kim, S. Lee, J. Lin, Z. Dujmic, W. Kim, J. Talag, A. Zuccolo, C. Fan, A. Sebastian, M. Kramer, L. Spiegel, L. Nascimento, T. Zutavern, B. Miller, C. Ambroise, S. Muller, W. Spooner, A. Narechania, L. Ren, S. Wei, S. Kumari, B. Faga, M. J. Levy, L. McMahan, P. Van Buren, M. W. Vaughn, K. Ying, C. T. Yeh, S. J. Emrich, Y. Jia, A. Kalyanaraman, A. P. Hsia, W. B. Barbazuk, R. S. Baucom, T. P. Brutnell, N. C. Carpita, C. Chaparro, J. M. Chia, J. M. Deragon, J. C. Estill, Y. Fu, J. A. Jeddeloh, Y. Han, H. Lee, P. Li, D. R. Lisch, S. Liu, Z. Liu, D. H. Nagel, M. C. McCann, P. SanMiguel, A. M. Myers, D. Nettleton, J. Nguyen, B. W. Penning, L. Ponnala, K. L. Schneider, D. C. Schwartz, A. Sharma, C. Soderlund, N. M. Springer, Q. Sun, H. Wang, M. Waterman, R. Westerman, T. K. Wolfgruber, L. Yang, Y. Yu, L. Zhang, S. Zhou, Q. Zhu, J. L. Bennetzen, R. K. Dawe, J. Jiang, N. Jiang, G. G. Presting, S. R. Wessler, S. Aluru, R. A. Martienssen, S. W. Clifton, W. R. McCombie, R. A. Wing and R. K. Wilson (2009). “The B73 maize genome: complexity, diversity, and dynamics.” Sci 326(5956): 1112–1115.doi: 10.1126/science.1178534.

Schneider, C. A., W. S. Rasband and K. W. Eliceiri (2012). “NIH Image to ImageJ: 25 years of image analysis.” Nat Methods 9(7): 671–675.

Somerville, C., H. Youngs, C. Taylor, S. C. Davis and S. P. Long (2010). “Feedstocks for lignocellulosic biofuels.” Sci 329(5993): 790–792.doi: 10.1126/science.1189268.

Stewart, C. N., Jr. and L. E. Via (1993). “A rapid CTAB DNA isolation technique useful for RAPD fingerprinting and other PCR applications.” Biotechniques 14(5): 748–750.

Thorvaldsdóttir, H., J. T. Robinson and J. P. Mesirov (2012). “Integrative Genomics Viewer (IGV): high-performance genomics data visualization and exploration.” Briefings in Bioinformatics 14(2): 178–192.doi: 10.1093/bib/bbs017.

Vicient, C. M., M. J. Jaaskelainen, R. Kalendar and A. H. Schulman (2001). “Active retrotransposons are a common feature of grass genomes.” Plant Physiol 125(3): 1283–1292.

Vicient, C. M., A. Suoniemi, K. Anamthawat-Jonsson, J. Tanskanen, A. Beharav, E. Nevo and A. H. Schulman (1999). “Retrotransposon BARE-1 and Its Role in Genome Evolution in the Genus Hordeum.” Plant Cell 11(9): 1769–1784.

Wang, S., N. Liu, K. Peng and Q. Zhang (1999). “The distribution and copy number of copia-like retrotransposons in rice (Oryza sativa L.) and their implications in the organization and evolution of the rice genome.” Proc Natl Acad Sci U S A 96(12): 6824–6828.

Wicker, T., F. Sabot, A. Hua-Van, J. L. Bennetzen, P. Capy, B. Chalhoub, A. Flavell, P. Leroy, M. Morgante, O. Panaud, E. Paux, P. SanMiguel and A. H. Schulman (2007). “A unified classification system for eukaryotic transposable elements.” Nat Rev Genet 8(12): 973–982.doi: 10.1038/nrg2165.

Woodard, K. R. and G. M. Prine (1993). “Dry-Matter Accumulation of Elephantgrass, Energycane, and Elephantmillet in a Subtropical Climate.” Crop Sci 33(4): 818–824.doi: DOI 10.2135/cropsci1993.0011183X003300040038x.

Wu, J., Y. Huang, Y. Lin, C. Fu, S. Liu, Z. Deng, Q. Li, Z. Huang, R. Chen and M. Zhang (2014). “Unexpected Inheritance Pattern of Erianthus arundinaceus Chromosomes in the Intergeneric Progeny between Saccharum spp. and Erianthus arundinaceus.” PLOS ONE 9(10): e110390.doi: 10.1371/journal.pone.0110390.

Xie, X. M., X. Q. Zhang, Z. X. Dong and H. R. Guo (2011). “Dynamic changes of lignin contents of MT-1 elephant grass and its closely related cultivars.” Biomass Bioenerg 35(5): 1732–1738.doi: 10.1016/j.biombioe.2011.01.018.

Xu, Y. and J. Du (2014). “Young but not relatively old retrotransposons are preferentially located in gene-rich euchromatic regions in tomato (Solanum lycopersicum) plants.” Plant J 80(4): 582–591.

Zedek, F., J. Smerda, P. Smarda and P. Bures (2010). “Correlated evolution of LTR retrotransposons and genome size in the genus Eleocharis.” BMC Plant Biol 10(1): 265.doi: 10.1186/1471-2229-10-265.

Zhang, L., X. Yang, L. Tian, L. Chen and W. Yu (2016). “Identification of peanut (Arachis hypogaea) chromosomes using a fluorescence in situ hybridization system reveals multiple hybridization events during tetraploid peanut formation.” New Phytologist 211(4): 1424–1439.

Zhao, L. J. (2010). “Karyotype Analysis and Ribosomal DNA Fluorescent in Situ Hybridization in Pennisetum purpureum.” Journal of Anhui Agricultural Sciences 38(22): 12170–12173.doi: 10.3969/j.issn.0517-6611.2010.22.193.

Zhu, X. G., S. P. Long and D. R. Ort (2008). “What is the maximum efficiency with which photosynthesis can convert solar energy into biomass?” Current Opinion in Biotechnology 19(2): 153–159.

