## Supplemental Table S1 for "Diversity chromosome evolution of Ty1-copia Retrotransposons in *Pennisetum purpureum* Revealed by FISH"

Table S1. Classification of PCR products into families based on amino acid similarity

| Element family | Serial number clones | Amino acid similarity | Stop coding or Frameshift mutations |
| --- | --- | --- | --- |
| Pc1 | 80,42,70,33,62,11,59,30,10,5,71,34,83,76,16,14,12,36,63 | 78.30 %-100 % | 1 |
| Pc2 | 29,20 | 83.78% | 2 |
| Pc3 | 52,74,13,61,53,28,79,38,49,68 | 79.70 %-100% | 1 |
| Pc4 | 6,24,58,8,39,25,48,41,37,50 | 81.0 %-100% | 2 |
| Pc5 | 9 | N/A | 1 |
| Pc6 | 26 | N/A | 1 |
| Pc7 | 56 | N/A | 1 |
| Pc8 | 3,64,21,66 | 93.4 %-100 % | 1 |
| Pc9 | 81 | N/A | 0 |
| Pc10 | 65 | N/A | 1 |
| Pc11 | 60 | N/A | 1 |
| Pc12 | 57 | N/A | 0 |
| Pc13 | 45,78 | 92.21% | 1 |
| Pc14 | 22 | N/A | 0 |
| Pc15 | 87 | N/A | 0 |
| Pc16 | 17 | N/A | 1 |
| Pc17 | 88,4 | 92.11% | 1 |
| Pc18 | 51 | N/A | 0 |
| Pc19 | 85,75 | 100% | 0 |
| Pc20 | 27 | N/A | 0 |
| Pc21 | 47,15 | 100% | 0 |
| Pc22 | 69,35 | 84.78% | 2 |
| Pc23 | 72,73 | 84.78% | 2 |
| Pc24 | 54 | N/A | 0 |
| Pc25 | 67,77 | 72.37% | 1 |
| Pc26 | 7 | N/A | 1 |
| Pc27 | 82,44 | 69.74% | 0 |
| Pc28 | 32,1,55,40,23,2,46,86,31,18,43,84,19 | 61.8 %-100 % | 3 |
